# Horizontal saccade bias results from combination of saliency anisotropies and egocentric biases

**DOI:** 10.1101/2025.04.21.648827

**Authors:** Stephanie M. Reeves, Jorge Otero-Millan

## Abstract

Saccadic eye movements shift the fovea between objects of interest to build a visual percept. In humans, saccades are predominantly executed along the cardinal axes, particularly in the horizontal direction. It is unknown how this horizontal saccade bias could arise mechanistically, though previous work suggests contributions from neural, image-based, and ocular motor factors. Here we used two publicly available eye movement datasets to first investigate which image features–spatial frequency, saliency, and structural content–relate to the horizontal saccade bias. Among the three image features, we found that orientation anisotropies in saliency content best predicted the strength of the horizontal saccade bias. Based on this result, we next implemented a saccade target selection model combining allocentric biases aligned with image orientation and egocentric biases aligned with eye or head orientation, independent of image content. As in prior work, this combination successfully replicated human saccade distributions during free viewing of upright images. When applied to tilted images, the model produced effects of image tilt and saccade size that were correlated with prior empirical findings, though with reduced amplitude, suggesting that current saliency models do not fully capture image effects. Taken together, these results suggest that saccade generation reflects both the allocentric biases present in the structure of natural scenes and the egocentric biases present in the saccade generation system itself. An open question is why the egocentric saccade bias exists, but our results suggest that it is adaptive in response to regularities in the world and our typical upright orientation.

## 1. Introduction

Although saccades can be executed in any direction, in humans, they occur more often in the cardinal directions than in oblique directions and in the horizontal direction more than in the vertical direction. This ocular motor anisotropy, which we will refer to as horizontal saccade bias, has been noted in a variety of settings and tasks including free viewing^1–7^, visual search^8,9^, driving^10^, navigating^10^, and during attempted fixation^3,11–14^. It is hypothesized that the horizontal saccade bias originates from a combination of ocular motor, neural, environmental, and learned biases, though the exact source is unknown^1,2,15,16^.

Prior work has confirmed that the natural environment has statistical regularities and is highly structured: objects tend to either lay on the ground due to gravity or stand straight up to fight gravity, resulting in a predominance of horizonal and vertical contours. This leads to visual scenes with high spectral power at cardinal orientations, and salient objects that are aligned vertically or horizontally^17–19^. These environmental regularities are thought to be related to certain perceptual anisotropies such as the oblique effect, which describes increased sensitivity to horizontal and vertical orientations over oblique ones, and has been documented in measures of visual acuity, contrast sensitivity, orientation discrimination, and recognition rate^20–22^. This is also manifested by anisotropies in filling-in at the blind spot ^23,24^. Previous work suggests that these perceptual anisotropies cannot be explained by the optics of the eye alone^25^, but rather from the overrepresentation of horizontal and vertical orientation selectivity in primary visual cortex^26–29^. Perception across the visual field is also not isotropic, as better visual performance has been documented along the horizontal than vertical meridian (horizontal-vertical anisotropy, HVA) and along the lower than upper meridian (vertical meridian asymmetry, VMA)^30^. Early theoretical and experimental work has shown that sensory neurons are adapted to environmental statistical regularities through a process of efficient coding^31,32^, and that environmental regularities play a role in biased neural representations in the brain. Recently, research has shown that neural biases in orientation may be instantiated in the brain through uneven tuning preferences across orientation, shedding light on how environmental statistics inform biological priors that produce perceptual anisotropies^26^.

To understand how the horizontal saccade bias could arise mechanistically, we consider how it may relate to other known orientation-related anisotropies. On one extreme, the saccade direction bias could be a complete result of orientation biases present in image content alone, with a neural circuitry for saccade generation that is isotropic across directions. In this case, viewing a stimulus without orientation cues would produce an isotropic saccade direction distribution. However, we know from previous work that the saccade direction bias is present even when fixating a circular dot in the dark^3,12–14^ and even when free-viewing a radially-symmetric image^4,33^. On the other extreme, the saccade direction bias could be a complete result of biases along the saccade generation pathway, including the visual system. In this case, because all those biases would be egocentric, any change in image orientation relative to the head should not influence saccade directions. However, prior work has found that the orientation at which an image is viewed influences saccade directions^2,4,33^, which suggests that biases in neural tuning in egocentric maps cannot be solely responsible for anisotropic saccade direction distributions.

Importantly, when individuals view Earth-upright natural scenes while their heads are tilted, we find that the saccade directions are neither oriented perfectly with respect to the scene nor with respect to the head: instead, the horizontal saccade bias is oriented somewhere in between the head and image reference frames^33^. These experimental data have led us to hypothesize that there may be a baseline, egocentric, ocular motor horizontal bias that humans are born with and develop^15^, and another bias that is stimulus-driven arising from the orientation biases present in the scene being explored at any given moment. It is yet unclear which scene properties may drive this image-based bias and how an image-based bias may interact with a lower-level egocentric ocular motor bias. The present work thus sought to determine: 1) which image features best predict the horizontal saccade bias, and 2) whether a model of saccade generation that combines a stimulus-driven image bias with a fixed egocentric saccade targeting bias can explain saccade direction biases in response to image tilt or head tilt.

## 2. Methods

### 2.1 Participants

We obtained human eye movement data that was recorded while participants viewed natural images. These data were taken from two different publicly available datasets: the DOVES dataset^34^ and the FVTilt dataset^11^ from the Ocular Motor lab. There were 28 unique participants from the DOVES dataset and 20 unique participants from the FVTilt dataset, totaling 48 participants. All participants provided informed consent before data collection and the research followed the tenets of the Declaration of Helsinki.

### 2.2 Stimuli

There were 101 natural images from the DOVES dataset and 40 images from the FVTilt dataset, totaling 141 images.

The DOVES images were obtained from a calibrated greyscale natural image database^26^ and were displayed on an Image Systems 21” greyscale gamma corrected monitor. The screen resolution was set at 60 pixels per 1° visual angle and the image display was about 17° x 13° visual angle. The FVTilt images were obtained from the MIT CSAIL database and were displayed on a 4K OLED screen. The screen resolution was 40 pixels per 1° visual angle and the image display was about 20° x 20° visual angle.

### 2.3 Procedure

All participants were seated and using a chin rest and/or bite bar to restrict head movement. All participants were instructed to freely view each image as it appeared, either for 5 s (DOVES) or 10 s (FVTilt).

Before each DOVES image appeared, the screen was filled with Gaussian white noise. Following stimulus presentation, participants were shown a small image patch (1° x 1°) and asked whether they had seen that image patch in the larger image shown previously. This memory task was used to keep observers motivated to explore the image.

Before each FVTilt image appeared, participants were instructed to fixate a central 0.5° dot. The full FVTilt dataset included some trials when the task of the participant was to fixate a dot with the image in the background or to free view an image could be tilted by -30° or 30°.

### 2.4 Eye movement data

As participants viewed each image, their eye movements were recorded using either a dual-Purkinje eye tracker (DOVES: monocular, 200 Hz, horizontal and vertical eye position data) or a custom-built video-based pupil eye tracker (OpenIris^35^) (FVTilt: binocular, 250 Hz, horizontal, vertical, and torsional eye position data). Both datasets used calibration techniques to match the output of the eye tracker (pixels) to position of the observer’s gaze in degrees of visual angle. Calibration occurred every 10 trials for DOVES and every 20 trials for FVTilt.

The raw eye movement samples were obtained from both datasets. Eye movement samples were first cleaned to remove blinks and other artifacts. Saccades were detected using a modified version of the algorithm designed by Engbert and Kliegl^36^ that is based on the robust standard deviation of eye movement data (λ=8 for FVTilt, λ=15 for DOVES). We reviewed saccade detection with visualizations of individual observer eye position traces (Figure 1A and B) in addition to summary plots of the main sequence (Figure 1C), amplitude distributions (Figure 1D), and saccade direction (Figure 1E).

**Figure 1.**
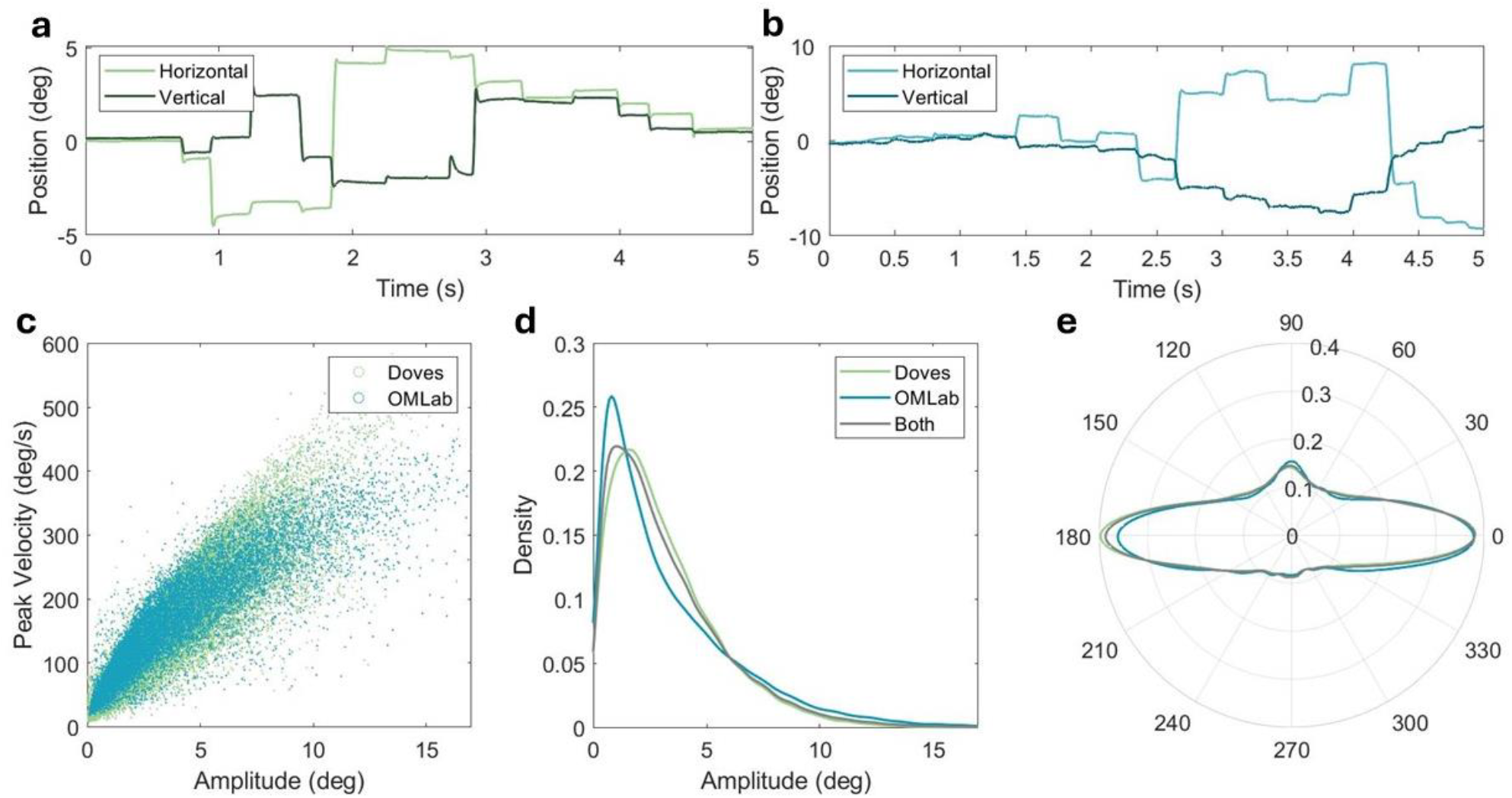
Raw eye movement and saccade statistics for both datasets. A) Horizontal and vertical eye movement traces from DOVES and B) FVTilt datasets. C) Main sequence for all saccades across both datasets. D) Amplitude distribution for all saccades across both datasets. E) Saccade direction distribution for all saccades across both datasets. Radial axis indicates probability and angle indicates saccade direction with 0° angle indicating saccade to the right and 180° angle indicating saccade to the left.

### 2.5 Image analysis

All images across the two datasets were analyzed for orientation anisotropies according to their spatial frequency, saliency, and structural content. Depiction of the image analysis pipeline is shown in Figure 2 using an exemplar image (Figure 2A).

**Figure 2.**
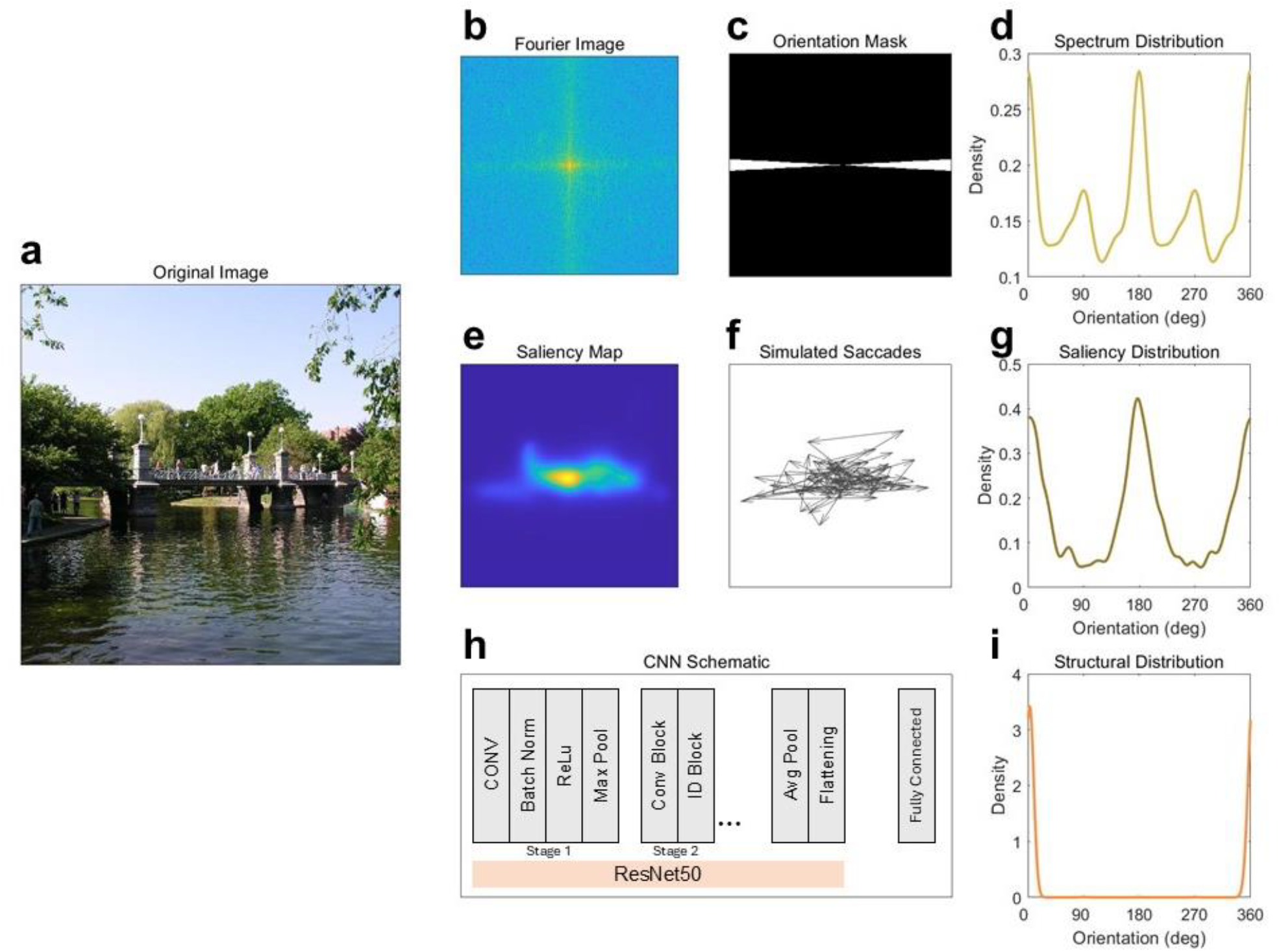
Depiction of image analysis pipeline for spatial frequency, saliency, and structural content. A) An exemplar input image. B) Fourier transform of the input image. C) Exemplar wedge orientation mask (scaled for visualization purposes). E) Saliency probability map of input image from DeepGaze IIE. F) Exemplar simulated saccades according to the saliency probability map. H) Illustration of CNN model that uses ResNet50 as the base model with a fully connected layer at the end. D, G, I) Density distribution as a function of orientation (0° and 180° indicate horizontal; 90° and 270° indicate vertical) according to each calculated image feature.

To analyze images according to their spatial frequency content, each image was first converted to greyscale and pixel values were scaled to range from 0 to 1. The image was multiplied with a hanning window to eliminate edge effects and then the mean pixel value was subtracted from all the pixel values to remove the DC component. We obtained the Fourier transform of the image (Figure 2B) and then used ring and wedge masks (Figure 2C) to calculate the amplitude spectrum as a function of spatial frequency and orientation. There were 10 spatial frequency bands split up in logarithmic steps ranging from 0.3 to 12.1 cycles per degree for the FVTilt dataset and ranging from 0.5 to 18.1 cycles per degree for the DOVES dataset. There were 180 orientation wedges (one wedge per degree with each wedge covering 10°) for both datasets. We calculated the amplitude spectrum as a function of orientation for each image by averaging across spatial frequency bands.

To analyze images according to their saliency content, we first obtained saliency prediction maps of each image using DeepGaze IIE^37^, a pretrained saliency prediction model that takes in an image and outputs a heatmap of probable fixation locations using low-level features and a center bias prior (Figure 2E). We used this saliency model as it does not include an explicit saccade direction bias (contains only a center bias) so that we could extract exclusively the biases present in the images. Other saliency models include an explicit not-image driven saccade direction bias^38^ or are trained with ordered saccade data must include the saccade direction bias^39,40^. We then simulated 2000 saccades according to that image’s saliency probability map (Figure 2F). We calculated the amplitude and direction of each simulated saccade and obtained a distribution as a function of angle.

To analyze images according to their structural content, we used a convolutional neural network (CNN) that was trained to determine the orientation of a given image relative to upright^41^ (Figure 2H). The model uses the pretrained ResNet50 as its base model and adds a fully connected layer to classify the images according to their orientation relative to gravity. This classifier was trained on Google Street View images and provides a 360-element vector containing the likelihood of each possible orientation angle of the image.

The three methods resulted in distributions as a function of orientation angle that we fit with a circular kernel density estimate (KDE) to obtain a smoothed distribution (Figures 2D, 2G, 2I).

### 2.6 Data analysis and statistics

For each image, we calculated a metric of horizontal orientation bias for: 1) the saccade behavioral data, and 2) each of the three image features. We obtained this metric of horizontal bias by fitting a given distribution with a circular KDE whose Gaussian bandwidth (kernel) was 0.1 radians. We averaged the density values at 0° and 180° to obtain a horizontal bias metrics for each image. To put the horizontal bias metrics into perspective, a simulated saccade distribution with only horizontal saccades (exclusively 0° and 180° directions) would produce a saccade horizontal bias metric of 2 using our chosen bandwidth of 0.1; and a simulated saccade distribution with uniform probabilities would produce a saccade horizontal bias metric of 0.16. We used a constant bandwidth across all analyses.

To determine whether any of the three image features could predict a change in the saccade horizontal bias, we used a linear regression model with one outcome variable and three predictor variables. We chose an alpha of 0.05 to determine significance.

### 2.7 Modeling saccade targeting

Saccade likelihood for each position in the scene (x,y) was calculated as a weighted combination of the likelihood given by the egocentric biases and the allocentric likelihood given by the content of the scene.

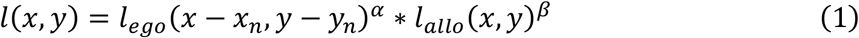

Where the allocentric likelihood is independent of the current position of the eye and the egocentric likelihood is always shifted by the current eye position. The allocentric likelihood was directly the output of the DeepGaze IIE saliency model and the egocentric likelihood was the result of combining a central bias *f*(*x, y*), a horizontal bias *v*(*x, y*), and a penalty for very small saccades *s*(*x, y*). With *x*^′^ = *x* − *x*_*n*_ and *y*^′^ = *y* − *y*_*n*_

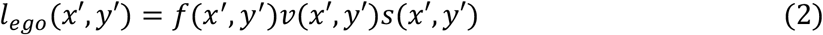

We found the parameters of the egocentric likelihood by maximizing the likelihood of the saccades present in the data from free viewing of upright images (Table S1). These parameters describe the egocentric biases derived from known properties of the human ocular motor system including the bias for small saccades over big saccades, modeled as a Cauchy distribution, and the bias for horizontal and vertical saccades over oblique saccades, modeled as a mixture of Von Mises distributions. We fit one parameter with the Cauchy distribution, *a* in equation 1, which describes a probability distribution for saccade endpoints (x’,y’).

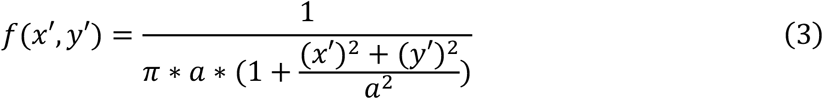

and twelve total parameters with von mises distributions (four lobes that each had a µ (circular mean), κ (circular standard deviation), and *w* (weight):

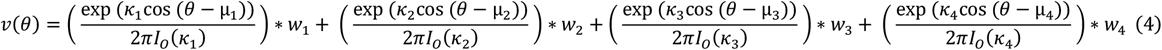

Here, *θ* = tan^−1^ (*y*^′^, *x*′) represents the angle of the vector from the current fixation to a particular pixel in the image. The four lobe weights (*w*) were constrained to sum to 1. For the model fitting, we used 15% of the upright-image free viewing data available from both datasets. Importantly, these data were from images with the weakest orientation bias according to our saliency orientation bias metric, which was important for isolating an estimate of the egocentric bias (separate from the influence of the image). We additionally implement one extra function to penalize small saccade generation based on prior work^3^.

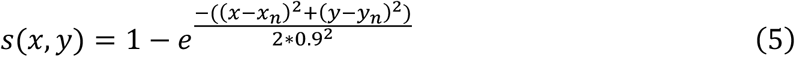

We fine-tuned our fitting procedures by simulating saccades according to combined probability maps comprised of weighted allocentric and egocentric maps. The weights (0.7 for egocentric map and 1.3 for allocentric map) were adjusted so as to find a good qualitative fit of both the saccade magnitude distributions and saccade direction distributions from saccades simulated with the model. Figure 3A-B shows an exemplar weighted allocentric saliency map (from one image) and the weighted egocentric map centered on the current saccade landing point. With each simulated saccade, the resultant combined map changes depending on the saccade location (Figure 3C). For this procedure, we used half of the upright images from the FVTilt dataset that were not used during the original fitting stage.

**Figure 3.**
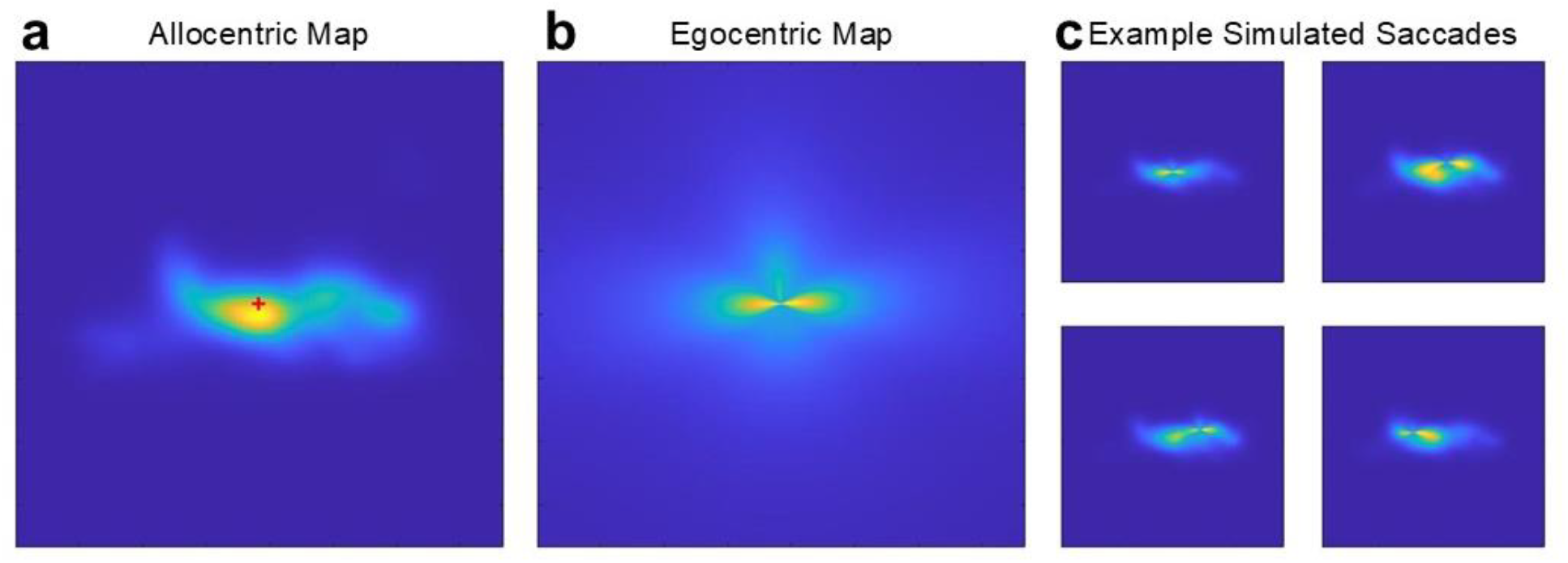
Example allocentric and egocentric saccade targeting maps. A) An allocentric probability map (saliency map) for an exemplar image. The red cross indicates the current saccade starting point. B) The egocentric probability map comprised of a direction mask and central bias mask centered on the saccade starting point. C) The resultant combined probability map for four consecutive saccades, obtained by multiplying the two allocentric and egocentric maps with some weight.

## 3. Results

### 3.1 Biases in saccade directions

To test whether different images produced different amounts of horizontal saccade bias, we fit eye movement saccade data for a given image, combined across subjects, with a circular kernel density estimate and took the average of the density estimate at 0° and 180° to calculate the horizontal bias metric. We found that the horizontal saccade bias was significantly modulated by the image itself with some images producing a strong horizontal saccade bias (Figure 4A, 4B) and other images producing a much weaker horizontal saccade bias (Figure 4C, 4D). We compared the variance of the horizontal bias metric with the variance of a metric due to chance (via shuffling image and eye movement data) and found a significant difference (F = 0.23, p<0.001, Figure 4E). This suggests that the strength of the saccade direction bias depends on the properties of the image itself and that it is not just the result of random variations.

**Figure 4.**
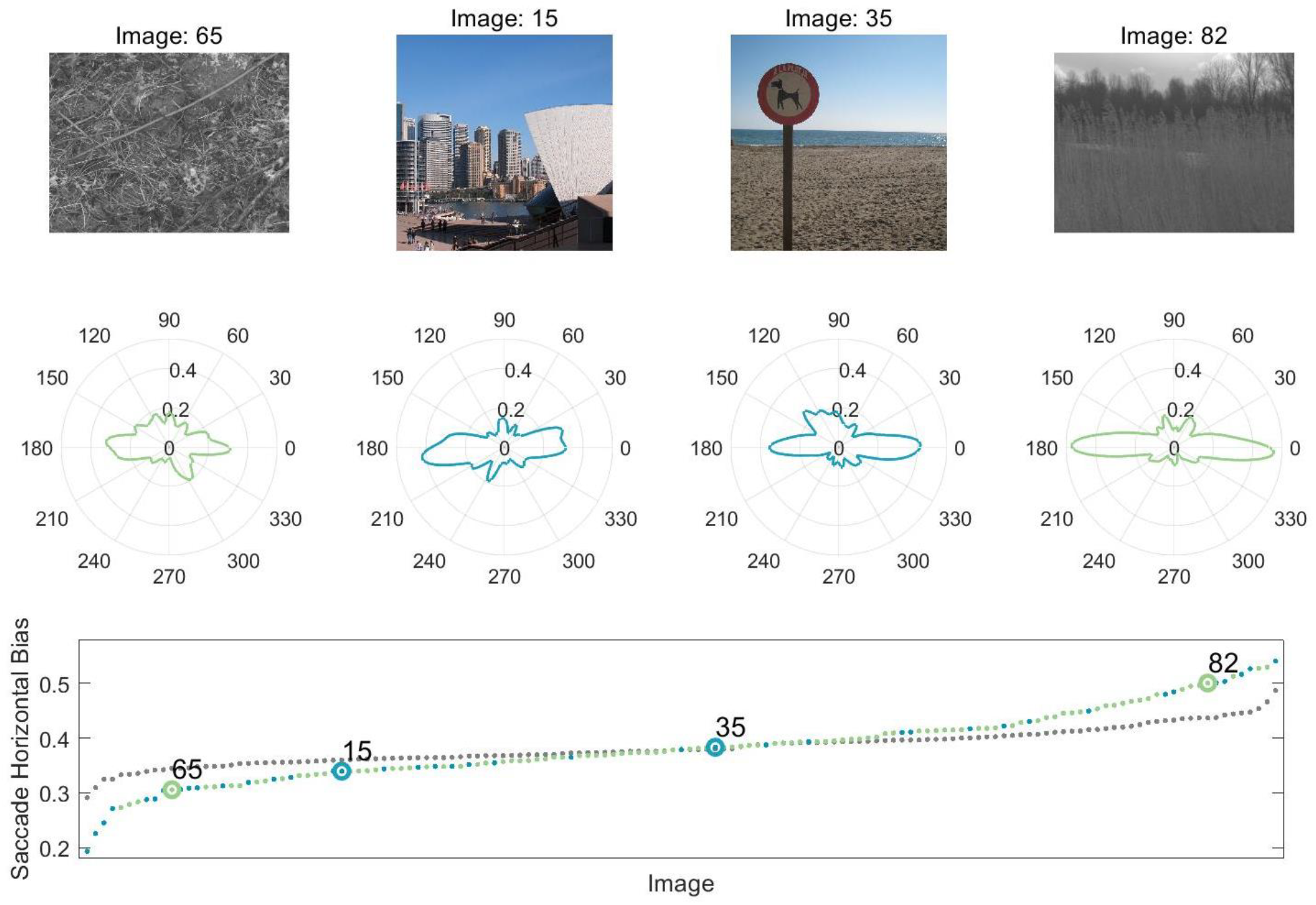
The saccade horizontal bias changes depending on the image. A) The saccade bias is strong for some images and D) weak for other images. B and C) Other images fall somewhere in between. E) Saccade horizontal biases change across images (green/blue dots) and differs from any difference we would have found due to chance (grey dots).

### 3.2 Saliency best predicts the strength of the horizontal saccade bias

Our main research question was whether any of the calculated image features would be predictive of the change in horizontal saccade biases across images. We implemented a linear regression model with one outcome variable, horizontal saccade bias, and three predictors, spatial frequency, saliency, and structural content. We found that the change in horizontal saccade biases across images could be significantly predicted by orientation biases in saliency content (β=0.36, p<0.001, Figure 5A). Spatial frequency and structural content were not predictive (p>0.65). The linear regression model is depicted in Figure 5B, and shows the incremental effect on the response of saliency caused by removing the effects of all other terms.

**Figure 5.**
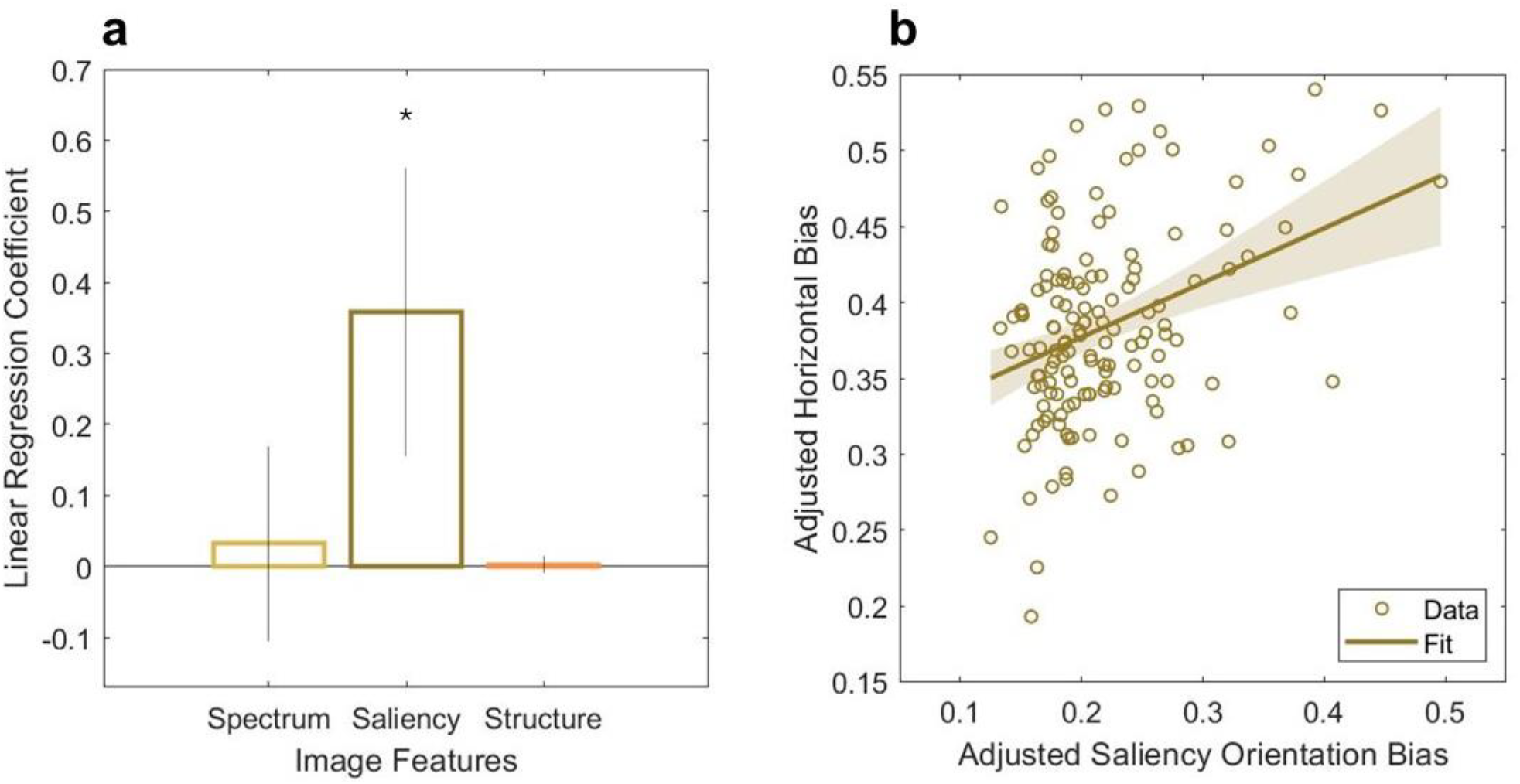
Linear regression model testing the influence of three image features on the horizontal saccade bias. A) Linear regression coefficients and 95% confidence intervals. B) Added variable plot, also known as a partial regression leverage plot, showing the influence of saliency on the horizontal bias from the linear model.

When analyzed separately, both datasets still result in saliency as the primary predictor for the change in the horizontal saccade bias (FVTilt: β=0.39, p=0.002; DOVES: β=0.50, p=0.005). This result suggests that strong orientation biases in the alignment of salient objects significantly predict the strength of saccade directional biases.

### 3.3 Comparing orientation biases between the two datasets

Obtaining two datasets for our analysis is advantageous because we can compare results between them. Orientation biases averaged across all images for the DOVES and FVTilt datasets are shown in Figure S3. We found predominant cardinal biases in spatial frequency content, horizontal biases in saliency content, and cardinal biases for structural content. To ensure our analyses were not biased due to edge effects or other artifacts, we computed orientation distributions for each image feature across 100 white noise images and observed relatively circularly uniform distributions across orientation per the Rayleigh’s test for non-uniformity of circular data (power spectrum: z<0.001, p=0.99; saliency: z<0.001, p=0.99; structural content: z=1.42, p=0.24).

### 3.4 Modeling saccade target selection

While orientation biases present in feature-based saliency maps are most predictive of the horizontal saccade bias, these saliency maps cannot fully account for the saccade direction bias. For example, we know that when humans view an Earth-upright natural scene while their head is tilted, saccade direction distributions do not completely align with the image and in fact are influenced by the orientation of both the head and scene^33^. To examine how a saccade direction bias may be instantiated in the brain, we implemented a model of saccade target selection that combined allocentric biases (aligned with image orientation) and egocentric biases (aligned with eye or head orientation). Our model receives an image as an input (upright or tilted) and then simulates a sequence of saccades by assigning each point in the image a likelihood to be selected as target, given the current fixation position (previous saccade endpoint) and a set of parameters to control for the amount the horizontal bias, central bias, and amplitude bias. See Methods for details.

Simulated saccade direction distributions in response to upright images are plotted in red in Figure 6A along with human saccade distributions in blue for ease of comparison. Like previous work^38^, we achieve good fits to upright image data with our full model. To compare our model against either an allocentric-only or egocentric-only strategy, we also simulated saccades according to an allocentric saliency map (green, Figure 6A) and according to an egocentric map (gold, Figure 6A). Our model generally produces saccade direction distributions that align with real behavioral data, unlike a saliency-only approach or an egocentric-only approach.

**Figure 6.**
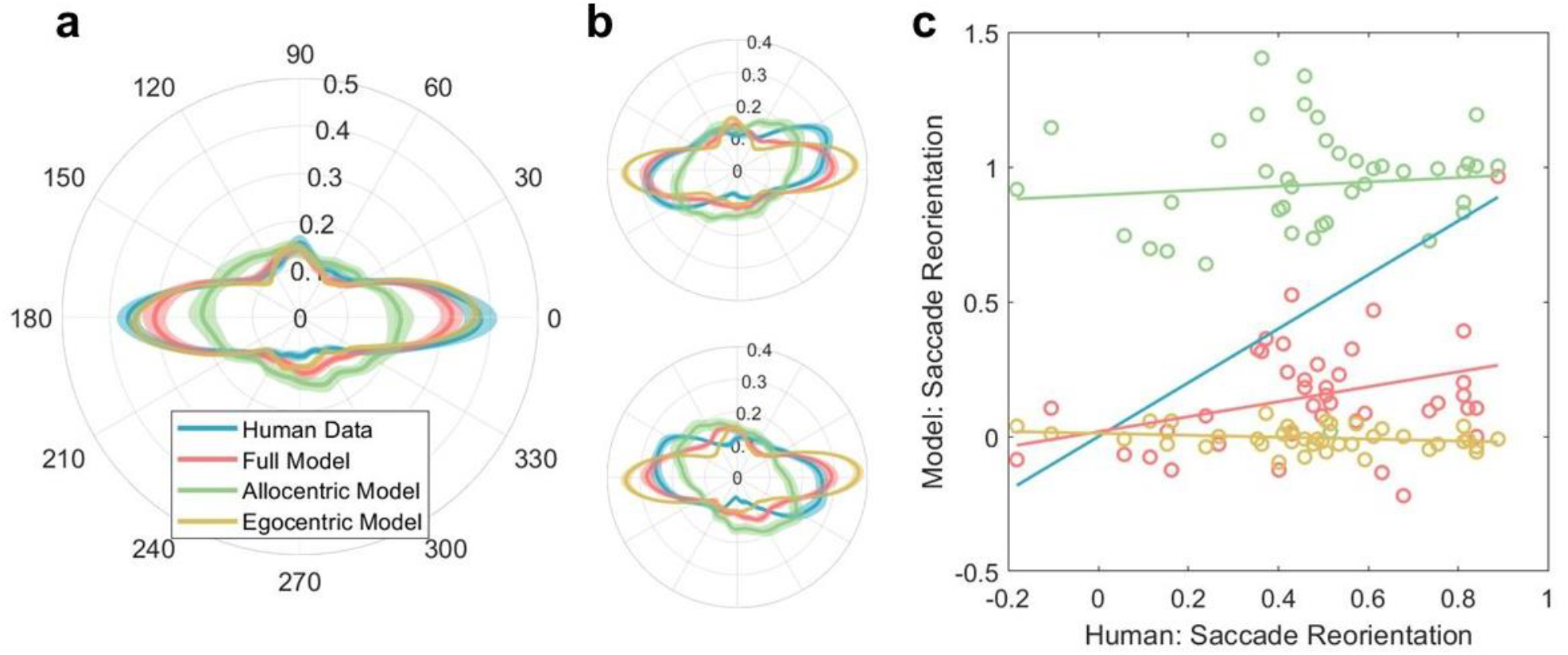
Human and simulated saccade direction distributions. A) Polar plot showing saccade direction distributions from human data (blue) and the full model (red) in response to an upright image using half the data (different from fitting). Green and gold data indicate the resultant distribution from simulating saccades according to an allocentric salience map alone and an egocentric map alone, respectively. B) Polar plot showing saccade directions in response to a -30° tilted image and 30° tilted image. C) Individual dots represent reorientation indices for individual images for both the human (x axis) and model data (y axis changes depending on which simulation is used) where 1 represents full reorientation and 0 represents no reorientation. Lines represent linear regression fits.

### 3.5 Model Prediction on Tilted Images

Previous work has already shown that saccade direction distributions rotate to partially align with image orientation when those images are tilted, termed “saccade reorientation.” To test whether our model could predict the reorientation of the saccade bias in response to a tilted image, we simulated saccades according to the model using tilted allocentric saliency maps combined with an egocentric bias. We calculated how well these simulated saccade distributions matched human saccade distributions by calculating the amount of reorientation compared to the 0° tilt distribution using cross correlation for each image^33^. These reorientation indices were then scaled such that 1 represented full reorientation in the direction of the image tilt while 0 represented no reorientation.

Figure 6B shows polar histograms comparing saccade direction distributions from human data and the model in response to image tilt. Although the saccade direction distributions predicted by the model are generally too horizontal, they more closely resemble the human data than either the allocentric-only (too isotropic) or egocentric-only (too horizontal) simulations. We quantified this by comparing reorientation indices—how much the saccade distribution rotates in response to image tilt—between model and human data, and found a significant correlation (rho = 0.33, p=0.03; Figure 6C). This indicates that images eliciting strong reorientation in humans also elicited strong reorientation in the model. In contrast, reorientation indices from the allocentric-only (rho = 0.09, p=0.58) and egocentric-only (rho = -0.21, p=0.18) simulations were uncorrelated with human data, as expected: allocentric-only simulations tend to over-rotate (indices near 1), while egocentric-only simulations show little to no rotation (indices near 0). Although the full model exhibited some reorientation in response to image tilt, the effect was weaker than in humans, suggesting that saliency alone cannot fully account for the horizontal bias and does not sufficiently rotate the saccade distribution.

### 3.6 Modeling the influence of saccade amplitude on saccade directions

Previous work has found that not all saccades respond the same to image tilt: large saccades are more likely to be executed with respect to the tiled natural scene, while small saccades are likely to be executed horizontally with respect to the upright head/body^8^. We wondered whether the simulated saccades would mimic this behavioral finding. To address this question, we partitioned the human saccade data into four groups according to saccade amplitude. Thresholds for this partitioning were set based on quartiles of the human data to ensure that each group contained the same number of saccades. Figure 7 shows the saccade direction distributions as a function of amplitude (Figure 7A-D) with the associated saccade bias reorientation indices. We found that the biggest saccades are most influenced by image orientation while the smallest saccades are least influenced by image orientation, supporting previous work^11^. When we fitted a linear mixed effects model to the relationship between amplitude and the saccade bias reorientation predicted by the model, we found that the saccade bias reorientation was significantly related to the saccade amplitudes (β = 0.09, 95% CI = [0.06, 0.12], p<0.001). The magnitude of the effect is not as large as behavioral data has shown previously^11^. The horizontal bias present in allocentric saliency images does not capture the horizontal bias fully and thus, while we find that our biggest simulated saccades reorient to align with the image, the overall reorientation is less than we would expect.

**Figure 7.**
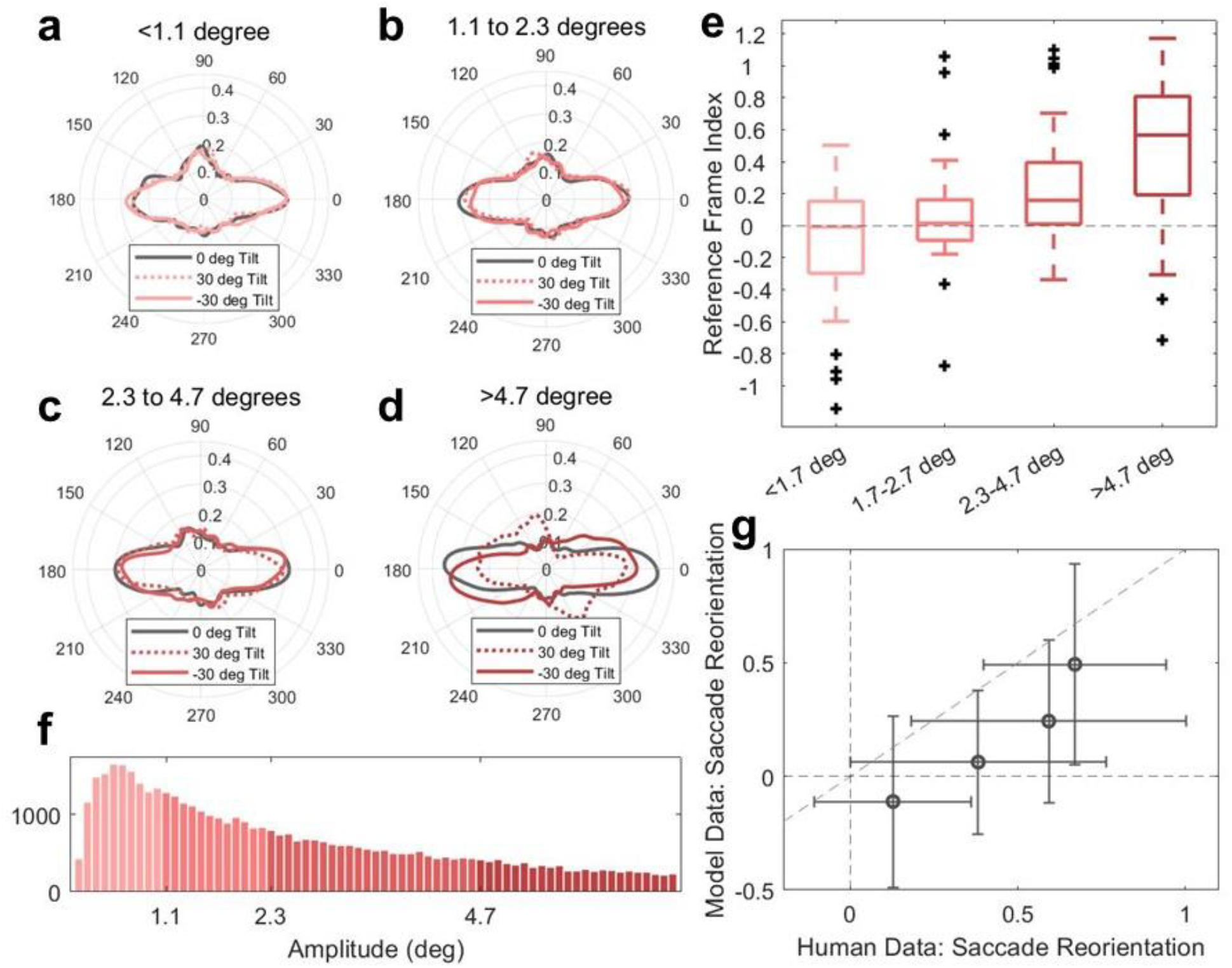
Model replication of amplitude-dependent saccade direction bias. A-D) Polar distributions of simulated saccade directions filtered by amplitude per our model. Amplitudes were determined based on quartiles of the data. E) Reference frame indices show that larger amplitude saccades were closely oriented to the tilt of the image (closer to 1) while smaller amplitude saccades were oriented egocentrically (closer to 0). F) Amplitude distribution of the simulated saccades. G) Dots represent the average reorientation indices across images for each of the four amplitude bins for both the human and model data. Error bars represent standard deviation; 1 represents full reorientation and 0 represents no reorientation.

## 4. Discussion

Humans make more saccades in the cardinal than oblique directions and more saccades in the horizontal than vertical direction. We know that this saccade direction bias is in part related to the current image being viewed because previous work has shown that saccade distributions rotate in response to the tilt of natural scenes^2^. The saccade bias is also related to the orientation of the head or retina since previous work has shown that the bias is orientated horizontally with respect to the head or eye during free viewing of a radially symmetric scene^33^ or during attempted fixation^3^. Here we sought to: 1) identify image features that may be predictive of a change in the saccade direction bias, and 2) describe the saccade direction bias in a model that combines inherent visual scene biases and egocentric ocular motor biases.

We found that images contain significant orientation biases in their spatial frequency, saliency, and structural content. Importantly, the extent of these orientation biases change depending on the image: a close-up image of leaves has less energy in cardinal orientations, weaker saliency, and fewer structural content clues than an image of a mountainous landscape. Of all the image-based orientation biases, we found that saliency was most predictive of changes in the horizontal saccade bias (Figure 5). Specifically, images that contained stronger orientation structure in saliency content resulted in saccade direction distributions that were more horizontal. However, we found that current saliency models failed to fully capture the influence of image tilt on saccade directions, as they accounted for only a fraction of its effect (to be discussed further below).

The use of two different datasets in these image-based analyses was integral because the content of the images across both datasets varied significantly: the FVTilt dataset contained colored images of streets, landscapes, buildings, people, homes, and food, while the DOVES dataset contained greyscale images of nature from different viewpoints. The differences in image content between the datasets resulted in different image-based statistics. Despite differences across the datasets, saliency was most predictive of the horizontal bias not only for the combined datasets but also for each dataset individually, validating our analysis approach.

### 5.1 Saliency Models

Although saliency is most predictive of the saccade direction bias, saliency alone cannot explain behavioral data because saccades are not executed in perfect alignment with image orientation. Recent saliency prediction models have acknowledged this limitation of an allocentric-only strategy of saccade selection and have used different approaches to bake in ocular motor biases to existing models (see ^39^ for current saliency model comparisons). For example, Le Meur and colleagues^38^ include an explicit horizontal saccade bias and then simulate saccades according to that bias combined with lower-level saliency maps. Others have used deep neural networks trained on real human data (that contain a horizontal saccade bias) to produce likely fixation locations within an image, successfully reproducing saccade direction biases in upright images^39,40^. However, none of these state-of-the-art saliency models have been systematically tested on tilted images to evaluate how well the saccade direction distributions from the model match behavioral data. Uncovering the conditions that make these saliency prediction models fail may help elucidate their limitations and ultimately improve them for biological plausibility.

### 5.2 Model

We presented a model that, given an input image in any orientation, simulates saccades based on a combination of saliency and egocentric oculomotor biases. When presented with an upright image, the model successfully generated saccade direction distributions that looked like human saccade distributions, consistent with previous findings^38^. When presented with a tilted image, the model produced saccades that were executed in the direction of image tilt but with a lower-than-expected gain (Figure 6B, 6C). This suggests that the horizontal bias embedded in current low-level saliency maps is insufficient to fully reorient simulated saccades in accordance with image tilt. However, saccade reorientation across images from the model was correlated with saccade reorientation across images from the human data, suggesting that the full model produced saccades in response to image tilt that were more similar to human saccades than either the allocentric-only or egocentric-only strategy.

When we tested whether our model could reproduce empirical findings that large saccades are likely to be executed with respect to a tilted image while small saccades are likely to be executed with respect to the head or retina, we found that the model broadly reproduced this finding (in the correct direction of the image tilt, Figure 7). However, the magnitude of this effect was smaller than what has been observed in human studies^11^. This again suggests that there is something about image saliency that we are not fully capturing.

### 5.3 Evolution/Development Across Species

The environment has played an active role in developing our ocular motor system over millennia. Orientation biases present in the environment likely drove humans to evolve with egocentric ocular motor priors in the same way they drove the evolution of perceptual and neuronal anisotropies. Thus, we may consider a phylogenetical egocentric ocular motor bias to be the cumulative result of repeated exposure to stereotyped environmental regularities. Over the lifespan, this phylogenetical ocular motor bias is fine-tuned and solidified developmentally^15^. As adults, the saccade direction bias can be flexibly modified depending on the current scene or visual experience to be more allocentrically-aligned or egocentrically-aligned depending on the task, body orientation, or size of saccade.

Other species, like macaques, live in different environments than humans and as a result, may show different egocentric ocular motor biases. Previous work has shown that macaques make predominantly vertical saccades during fixation^42–44^, but it is an open question what macaque saccade direction distributions may look like during a more natural free viewing task. It is reasonable to hypothesize that ocular motor biases present in other species are similarly a result of that species’ environmental regularities, and a comprehensive evaluation of saccade direction biases across species could shed light on some of these questions.

### 5.4 Egocentric Bias in the Brain

One open question is where this human egocentric saccade direction bias may be instantiated in the brain. This bias could theoretically occur in any of the three phases of saccade generation: target selection (involving the superior colliculus with inputs from FEF, parietal eye fields, supplementary eye fields, and visual areas), saccade execution (involving the burst generator in the brainstem), or saccade calibration (involving the cerebellum). If the bias was instantiated during target selection (i.e., within or before the superior colliculus), that could mean that there are simply more neurons preferring horizontal saccades than saccades of other directions. Early work by Robinson found that direct stimulation of the superior colliculus in monkeys evokes saccades with reliable directions and amplitudes that do not depend on the amount of stimulation or the initial eye position. The elicited saccades were predominantly horizontal with some small vertical components^45^. Stimulation of the superior colliculus in mice similarly elicited predominantly horizontal saccades^46^. This could indicate that an egocentric saccade direction bias resides in superior colliculus, though certainly other areas involved in saccade planning such as FEF are possible as well, with a recent study showing that most neurons in FEF prefer horizontal saccades^47^. Alternatively, if the bias was instantiated during saccade execution, that could suggest saccade landing errors (i.e., a selected target is not reached successfully). By far the most common type of saccade landing error reported in the literature is radial undershoots in response to a single target, with greatest accuracy for saccades directed horizontally or cardinally^48,49^. Lastly, if the bias was instantiated in the cerebellum, then when the cerebellum is impaired we might see a reduced horizontal saccade bias. Patients suffering from cerebellar disorders report decreased saccade velocities and saccade amplitudes^50^; to our knowledge, no documentation of saccade direction changes have been reported.

### 5.5 Conclusions

We sought out to identify image-related features that could predict the strength of the horizontal saccade bias and build a mechanistic model of saccade generation in attempts to explain saccade direction biases. Our findings indicate that saliency content best explains variations in the horizontal saccade bias, surpassing spatial frequency and structural content. However, saliency alone does not fully account for the observed horizontal bias in saccade distributions; simulations based solely on saliency produce saccade patterns that are not sufficiently horizontal. Incorporating an egocentric horizontal saccade bias into our model improved alignment with human behavioral data for upright images and partially reproduced human findings related to image tilt and saccade amplitude, though not with the expected gain. Testing the performance of the model in conditions of image tilt revealed that future improvements to saccade target prediction models may need to incorporate image tilt and head tilt to assess biological plausibility, as these conditions may reveal the model’s limitations and further refine its predictive power.

## 5. Data Availability

The eye movement data used in this manuscript have been previously published and made available for public download by van der Linde and colleagues^34^ and Reeves and colleagues^11^.

## 6. Acknowledgements

Funding for this work was provided by the National Eye Institute Award R00EY027846, the National Institutes of Health Training Grant 5T32EY007043-43, and the UC Berkeley Center for the Innovation in Vision and Optics (CIVO). The authors would like to thank Emily A Cooper for her comments.

## 8. Author Contributions

SMR and JOM designed the research, SMR conducted the data analysis, SMR and JOM directed and reviewed analysis, SMR and JOM wrote and reviewed the manuscript.

