## Supplementary Material for "Horizontal saccade bias results from combination of saliency anisotropies and egocentric biases"

This file includes:

Figure S1 to S3.

Table S1.

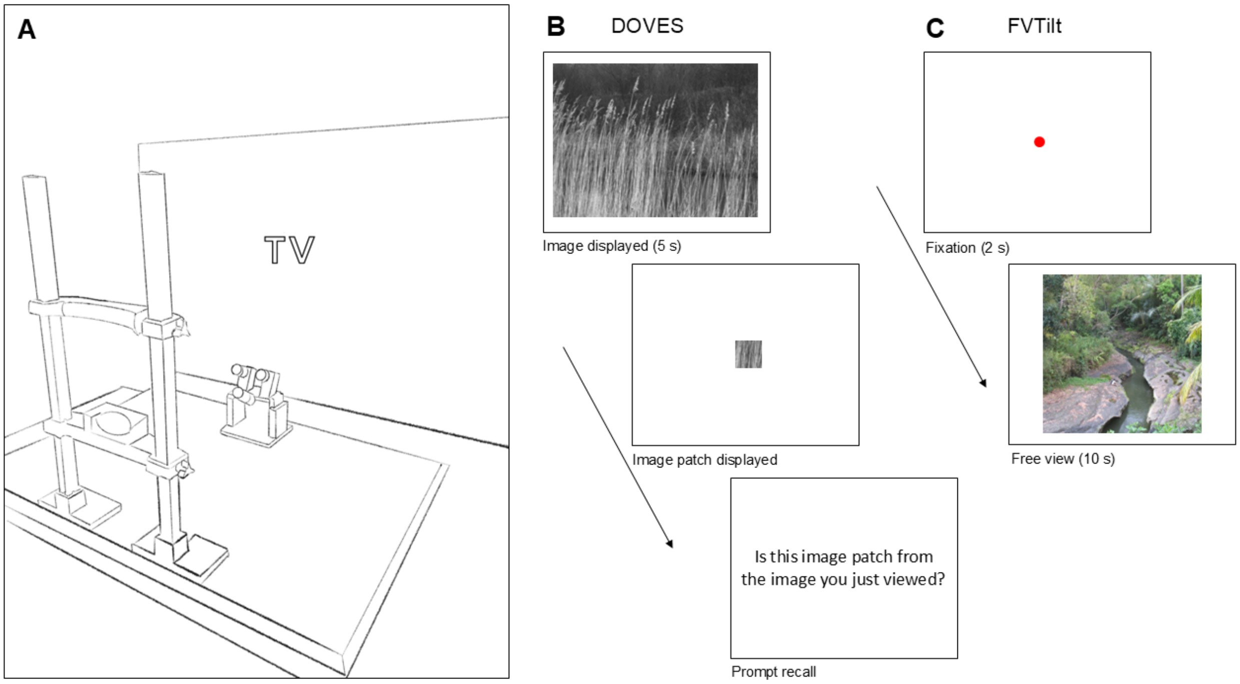


Figure S1. Experimental setup and procedure for the FVTilt and DOVES datasets. A) Experimental set-up for the FVTilt dataset. B) Illustration of the trial sequence for the DOVES dataset and C) FVTilt dataset.

Table S1. Description of model parameter estimates.

| **Parameter** | **Estimated Value** | **Description** |
|---|---|---|
| $a$ | 0.00002 | Cauchy distribution scale parameter |
| $\mu_1$ | -3.12 | Mean von mises lobe 1 |
| $\kappa_1$ | 13.36 | Standard deviation von mises lobe 1 |
| $w_1$ | 0.17 | Weight von mises lobe 1 |
| $\mu_2$ | 0.045 | Mean von mises lobe 2 |
| $\kappa_2$ | 16.13 | Standard deviation von mises lobe 1 |
| $w_2$ | 0.17 | Weight von mises lobe 2 |
| $\mu_3$ | 1.67 | Mean von mises lobe 3 |
| $\kappa_3$ | 11.55 | Standard deviation von mises lobe 3 |
| $w_3$ | 0.05 | Weight von mises lobe 3 |
| $\mu_4$ | -1.98 | Mean von mises lobe 4 |
| $\kappa_4$ | 0.0000004 | Standard deviation von mises lobe 4 |
| $w_4$ | 0.59 | Weight von mises lobe 4 |

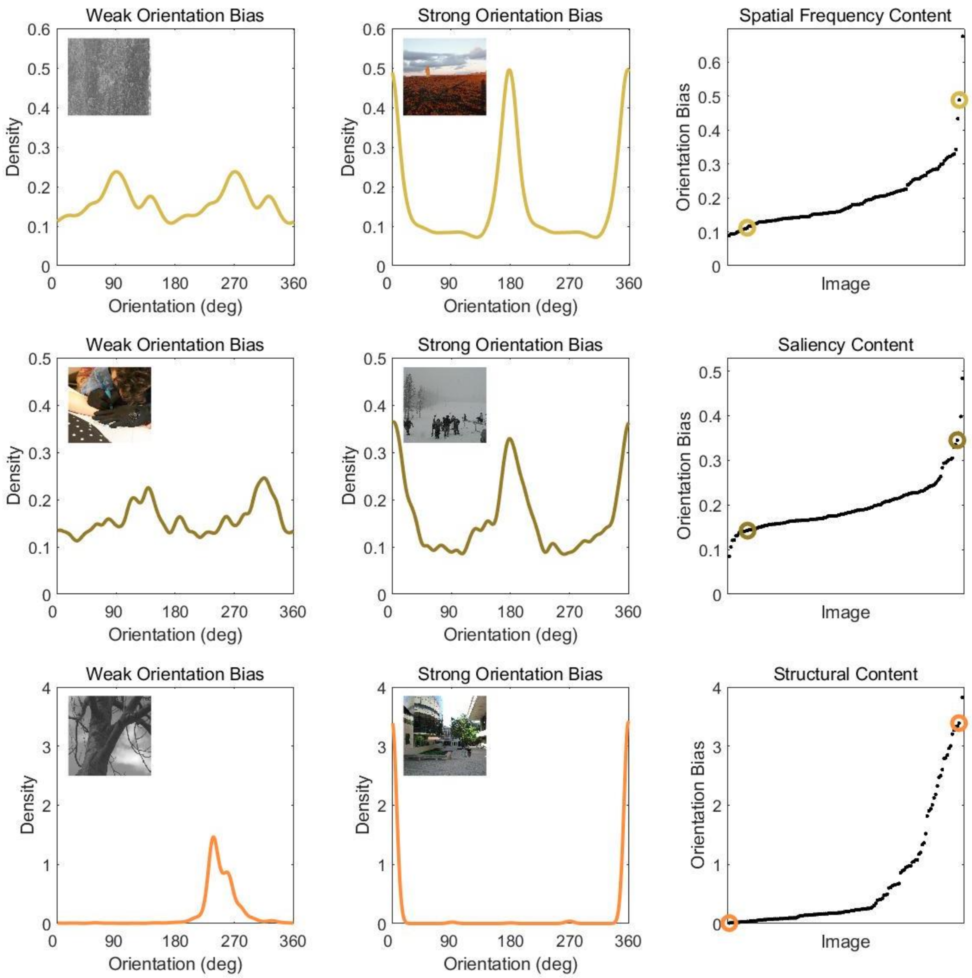


Figure S2. Results from image analyses for each of the three image features: A) spatial frequency content, B) saliency content, and C) structural content. The spatial frequency content density represents the relative spectral power along each direction (A), the saliency content density represents the likelihood of a saccade in a particular direction according to a saliency map (B), and the structural content density represents the estimated relative likelihood of the orientation of the image relative to gravity (C). Because the densities are normalized this gives an indication of the image's horizontal bias. For each image feature, an exemplar image containing weak (left) and strong (right) orientation biases are depicted. Angles 0° and 180° indicate horizontal; 90° and 270° indicate vertical. The right-most column shows the image-based orientation bias metric across images for that image feature. Note that x axis is sorted from low to high orientation bias within each image feature.

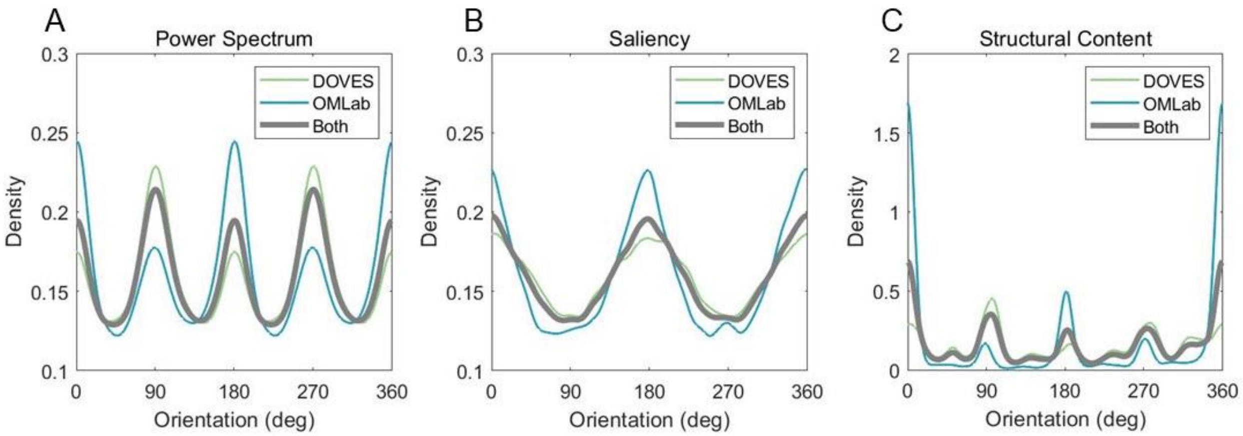


Figure S3. Depiction of orientation anisotropies present in the two datasets. Density distributions across orientation averaged across sets of images for A) spatial frequency content, B) saliency content, and C) structural content.
